# Hormone-sleep interactions predict cerebellar connectivity and behavior in aging females

**DOI:** 10.1101/2022.08.30.505858

**Authors:** Hannah K. Ballard, T. Bryan Jackson, Tracey H. Hicks, Sydney J. Cox, Abigail C. Miller, Ted Maldonado, Jessica A. Bernard

## Abstract

Sex hormones fluctuate over the course of the female lifespan and are associated with brain health and cognition. Thus, hormonal changes throughout female adulthood, and with menopause in particular, may contribute to sex differences in brain function and behavior. Further, sex hormones have been correlated with sleep patterns, which also exhibit sex-specific impacts on the brain and behavior. As such, the interplay between hormones and sleep may contribute to late-life brain and behavioral outcomes in females. Here, in a sample of healthy middle-aged and older females (n = 79, ages 35-86), we evaluated the effect of hormone-sleep interactions on cognitive and motor performance as well as cerebellar-frontal network connectivity. Salivary samples were used to measure 17*β*-estradiol, progesterone, and testosterone levels while overnight actigraphy was used to quantify sleep patterns. Cognitive behavior was quantified using the composite average of standardized scores on memory, processing speed, and attentional tasks, and motor behavior was indexed with sequence learning, balance, and dexterity tasks. We analyzed resting-state connectivity correlations for two specific cerebellar-frontal networks: a Crus I to dorsolateral prefrontal cortex network and a Lobule V to primary motor cortex network. In sum, results indicate that sex hormones and sleep patterns interact to predict cerebellar-frontal connectivity and behavior in aging females. Together, the current findings further highlight the potential consequences of endocrine aging in females and suggest that the link between sex hormones and sleep patterns may contribute, in part, to divergent outcomes between sexes in advanced age.

**Highlights:**

- Hormone-sleep interactions influence resting-state cerebellar-frontal connectivity
- Cognitive and motor behavior are impacted by hormone-sleep relationships
- Sex hormones and sleep patterns may contribute to aging outcomes in females

## 1. Introduction

Fluctuations in sex hormones throughout the female lifespan correlate with changes in volumetric and functional properties of the brain. Menopause is associated with grey matter reductions in the cerebellum (CBLM) and frontal cortex, and hormone therapy (HT) in postmenopausal females may reverse these changes (Rehbein et al., 2021). Hormone fluctuations during menstrual cycling modulate intrinsic connectivity in resting-state networks (Pritschet et al., 2020), and differences in functional connectivity emerge between reproductive and postmenopausal females (Ballard et al., 2022a). Further, brain changes during hormonal transitions are linked to attenuations in behavior as postmenopausal females undergoing HT exhibit improved cognition, compared to controls (LeBlanc et al., 2001).

Interestingly, changes in sleep patterns are tied to hormone fluctuations. Lower levels of sex hormones relate to increased insomnia, shorter deep sleep, and reductions in the physiological correlates of sleep to learning and memory consolidation (Brown and Gervais, 2020). Increases in sleep fragmentation and reductions in sleep quality occur with female reproductive aging as hormones decline (Brown and Gervais, 2020). In fact, sleep quality can be improved with HT in postmenopausal females (Li et al., 2015), whereas sleep fragmentation is correlated with increased risk of developing dementia (Sabia et al., 2021), which is more prevalent in females (Alzheimer’s Association, 2021). Thus, a balanced profile of sex hormones is necessary for maintaining proper sleep, and the interplay between these biological, age-linked factors may contribute to the brain and behavioral outcomes observed in female adulthood.

Relatedly, sleep patterns change with age in both females and males; further, sex-specific impacts of age-related sleep changes have been documented. Though both sexes display reduced sleep quality and increased sleep fragmentation with older age, the slope of these changes is significantly steeper in females (Li et al., 2021). In addition, the female brain appears to be especially vulnerable to the effects of sleep deprivation. After sleep deprivation, females exhibit greater reductions in resting-state connectivity and worsened cognition, compared to their male counterparts (Hajali et al., 2019). As such, sex differences in aging trajectories may be partially attributed to the interaction between hormone declines with menopause and modifications in sleep with age.

CBLM-frontal networks are susceptible to the effects of hormones and sleep. Our prior work revealed lower CBLM-frontal connectivity in postmenopausal females, relative to reproductive, including that between Crus I and the dorsolateral prefrontal cortex (DLPFC) as well as between Lobule V and the primary motor cortex (M1) (Ballard et al., 2022a). Crus I and the DLPFC play a role in cognitive function, while Lobule V and the M1 facilitate motor behaviors (O’Reilly et al., 2010). As such, we have shown that postmenopausal females display lower connectivity in both cognitive and motor-focused CBLM-frontal networks, which may have important implications for functional declines with age.

Work demonstrates that HT in postmenopausal females is associated with an increase in CBLM volume and improved cognition (Ghidoni et al., 2006). This corroborates the CBLM’s sensitivity to sex hormones, which is further supported by its abundance of estrogen receptors (Hedges et al., 2018). Estrogen also acts on the frontal cortex (e.g., DLPFC), which contains a substantial distribution of estrogen receptors as well (Hara et al., 2015). Thus, the frontal cortex is involved in estrogen signaling, in parallel with the CBLM, and these territories may be important with respect to endocrine aging.

CBLM-cortical interactions occur during sleep and may contribute to sleep-mediated memory consolidation (Canto et al., 2017). CBLM and frontal regions are modified by inadequate sleep and show sex differences in their relationship with sleep. In rats, sleep deprivation initiates a decrease in CBLM protein and glycogen levels (Gip et al., 2002). In humans, reductions in prefrontal activity are observed with sleep deprivation and correlate with impairments in cognition (Thomas et al., 2000). However, the role of CBLM-frontal networks in hormone-sleep interactions, and their contribution to differential aging outcomes between sexes, is relatively unexplored.

A robust body of literature suggests that females endure more severe consequences with age-related declines. In normative aging, females show steeper rates of cognitive impairment and motor deficiency than aging males (Levine et al., 2021). Anstey et al. (2021) revealed a shift in sex differences with rates of memory deterioration, where females convert to having steeper memory declines in older age compared to males, who have greater declines in middle-age. This shift may reflect impacts of the mid-life menopausal transition. Further, animal work reports steeper declines in memory in older females, compared to males, as the onset of some memory declines occur earlier in the female lifespan and coincide with changes in the estrous cycle (Markowska, 1999).

While several investigations note a relationship between hormone fluctuations and sleep changes throughout female adulthood, little work studies the role of these interrelated factors in aging outcomes for middle-aged (MA) and older adult (OA) females. These modifiable factors may offer potential therapeutic avenues for aging females as the benefits of HT and sleep intervention might be useful in rectifying the burdens of menopause. However, the contributions of CBLM-frontal networks in female-specific aging trajectories is, at present, widely unknown. Insight on the neural underpinnings of hormone-sleep interactions and their influence on behavior is necessary to understand sex discrepancies in functional declines.

To address these gaps, we conducted a cross-sectional study in healthy MA and OA females. We used salivary hormone measures and actigraphy-derived sleep assessments to predict cognitive and motor performance along with resting-state CBLM-frontal connectivity within two networks, as discovered in our prior work: Crus I-to-DLPFC and Lobule V-to-M1. We hypothesized that higher estrogen and progesterone levels and better sleep habits would jointly predict improved behavioral performance and greater resting-state connectivity across females, but especially in postmenopausal groups. We did not anticipate significant effects of testosterone, given the stability of this hormone across female adulthood.

## 2. Methods

Additional details regarding our methodology and full results are reported in **Appendix A** and **Appendix B**, respectively.

### 2.1 Participants

We included MA and OA females (ages 35-86) to compare across reproductive stages and examine the transition to menopause. We initially enrolled a total of 84 females in the study. Exclusion criteria were history of neurological disease, stroke, or formal diagnosis of psychiatric illness and use of contraceptives involving oral/intrauterine hormone administration or HT. These exclusions were made to evaluate impacts of normative endocrine aging on healthy adult females. One participant was lost to attrition and excluded from analyses. In addition, only three females were determined to be in perimenopause; as such, these participants were excluded from analyses. One late postmenopausal female was excluded from analyses due to suspected hormone use or a possible underlying endocrine condition. After excluding these participants, our initial sample consisted of 79 females (ages 35-86, mean age 59.15 ± 12.52). We did not exclude based off of handedness, thus right-handed (n = 77), left-handed (n = 0), and ambidextrous (n = 2) participants were included. All study procedures were approved by the Institutional Review Board at Texas A&M University, and written informed consent was attained from each participant.

### 2.2 Experimental Design

An overview of experimental procedures is illustrated in **Figure A1**. During the first session, participants provided saliva samples for hormone testing before performing a battery of cognitive/motor tasks. This was followed by an interim period of at least 2 weeks, during which participants completed questionnaires and wore an overnight actigraphy watch that gauged sleep patterns across 10 consecutive days. The time between sessions varied for some participants (n = 37), due to unexpected restrictions on data collection in response to the Covid-19 pandemic. On average, the interval between sessions was 37.68 ± 36.65 days across participants. Participants then returned for the second session, which included structural and resting-state magnetic resonance imaging (MRI). After concluding the study, participants completed a monthly menstrual tracking diary for 6 months to survey cycle characteristics.

### 2.3 Hormone Testing

Saliva samples were collected in pre-labeled cryovials provided by Salimetrics using the passive drool technique. Participants were asked to supply 1mL of saliva, after which samples were immediately stored in a -80 degrees Fahrenheit bio-freezer for stabilization. Assays were completed by Salimetrics to quantify estrogen (or 17*β*-estradiol), progesterone, and testosterone levels. The protocol used by Salimetrics includes two repetitions of each assay; thus, the values used in our analyses represent an average of both repetitions. A few samples were insufficient in quantity and were unable to be properly assayed (n = 3; 2 progesterone, 1 testosterone). The intra-assay coefficient of variability for our hormone samples was 0.15 for 17*β*-estradiol, 0.11 for progesterone, and 0.07 for testosterone.

Salivary measures have been strongly correlated with blood-derived measures for indexing sex hormone levels (17*β*-estradiol: *r* = 0.80; progesterone: *r* = 0.80; testosterone: *r* = 0.96, retrieved from https://salimetrics.com/analyte/salivary-estradiol/). The amount collected for our study is sufficient to detect 17*β*-estradiol at a high sensitivity threshold of 0.1 pg/mL (Salimetrics, 2006), along with 5.0 pg/mL and 1.0 pg/mL thresholds for progesterone and testosterone, respectively. Therefore, this non-invasive method is adequate for precisely measuring reproductive hormones.

### 2.4 Reproductive Staging

To categorize reproductive stage, we used responses on the menstrual tracking diaries, gathered over a period of 6 months, in tandem with the Stages of Reproductive Aging Workshop (STRAW+10) criteria (Harlow et al., 2012).

Participants with sufficient data were organized into 4 groups: reproductive, perimenopause, early postmenopause, and late postmenopause. Those with regular cycles, or variability of less than 7 days between consecutive cycles or across cycle lengths, were classified as reproductive. Females with 7+ days of variability between cycles or cycle lengths, 60+ days of amenorrhea, and/or ≤ 1 year since their final menstrual period (FMP) were determined to be in perimenopause. However, as only 3 females (ages 36, 42, and 56) were classified as perimenopausal using this approach, the perimenopause group was subsequently excluded from analyses. To distinguish between early and late postmenopausal females, we used the time, in years, since FMP. Those with 2-8 years since FMP were classified as early postmenopausal, and those with 9+ years since FMP were labeled as late postmenopausal. The resulting group characteristics are presented in **Table A1**.

Notably, there is observable age overlap across stage groups **Figure A2**, which indicates that our staging approach may have, in part, offset the inherent impact of age that is temporally associated with menopause.

### 2.5 Behavioral Testing

All participants completed a battery of cognitive and motor tasks to quantify behavior across these domains. To assess attention and executive function, we used the Stroop Task (Stroop, 1935). Performance was scored as the percent of correct responses on incongruent trials, including missed trials/those that were not responded to within the allotted time of 1.5 seconds. Participants that missed 90% or more trials were excluded from calculations as this suggests that they were not appropriately performing the task (n = 1; 100% of incongruent trials missed). Those with incomplete data were also excluded from calculations for this task (n = 1; 9/10 blocks incomplete). To index memory and semantic processing, we also included the Shopping List Memory Task modeled after Flegal et al. (2016). Scores for this task were calculated as the percentage of correct responses out of the 30 total trials, including misses. Finally, to gauge working memory and processing speed, participants performed the Digit-Symbol Substitution Task from the Wechsler Adult Intelligence Scale, 4^th^ Edition (WAIS-IV) (Benson et al., 2010). The total number of items completed (maximum of 135) was used to score performance on this task. Raw scores from each component of the cognitive battery (i.e., Stroop Task, Shopping List Memory Task, and Digit-Symbol Substitution Task) were first z-scored across the sample. These z-scores were then averaged together across all tasks to create a composite cognitive score for each participant.

For our motor variable, a Postural Sway Task was administered to measure properties of balance and stability. We calculated the 95% confidence interval (CI) of the center of pressure (COP) area (or sway area) using principal component analysis and a 9^th^ order Butterworth filter with a 20 Hz low-frequency threshold (Oliveira et al., 1996). The 95% CI of the COP area was then averaged across four separate positions for an overall score on the Postural Sway Task. To assess unimanual and bimanual dexterity, fine motor control, and coordination, we also administered the Purdue Pegboard Task (Tiffin and Asher, 1948). Scores for each test type were averaged across trials before summing all tests together for one comprehensive score on this task. Finally, we included an explicit Sequence Learning Task modeled after Kwak (Kwak et al., 2012) to evaluate motor learning and skill acquisition. To score this task, we calculated the percent of correct responses on the sequence trials (n = 324), including missed trials. Mirroring our approach for the cognitive tasks, scores for each motor task (i.e., Postural Sway, Purdue Pegboard, and Sequence Learning) were z-scored then averaged together to create a composite motor score.

### 2.6 Sleep Assessments

Sleep patterns were evaluated using ActiLife wGT3X-BT actigraphy watches, which produce results that are highly correlated with polysomnographic readings (Quante et al., 2018). Participants were asked to wear the actigraphy device on their non-dominant wrist for 10 consecutive days during the interim period between behavioral and imaging visits. We identified the average total sleep time across the 10-day period, which represents our sleep quantity variable. To quantify sleep quality, we extracted values for average sleep efficiency across all 10 days. Sleep periods were also corroborated with subjective sleep diaries completed during the interim.

### 2.7 Neuroimaging

During the second experimental session, participants underwent structural and resting-state MRI using a Siemens Magnetom Verio 3.0 Tesla scanner and a 32-channel head coil. Scanning protocols were adapted from the multiband sequences developed by the Human Connectome Project (HCP) (Harms et al., 2018) and the Center for Magnetic Resonance Research at the University of Minnesota. In total, participants underwent about 45 minutes of scanning, including a 1.5-minute localizer.

fMRIPrep (version 20.2.3, https://fmriprep.org/) was used to preprocess anatomical and functional images. The fMRIPrep preprocessing pipeline includes basic steps such as co-registration, normalization, unwarping, noise component extraction, segmentation, and skull-stripping. After denoising data using a band-pass filter of 0.008-0.09 Hz, first-level analyses were performed in Conn Toolbox, version 21a (Whitfield-Gabrieli and Nieto-Castanon, 2012), with a bivariate correlation approach. This work was supported in part by the Texas Virtual Data Library (ViDaL), funded by the Texas A&M University Research Development fund.

We included two CBLM regions of interest (ROIs) in our first-level analyses: Crus I and Lobule V. Crus I is associated with cognitive function whereas Lobule V is implicated in motor behavior (O’Reilly et al., 2010), and both regions are impacted in aging (Bernard and Seidler, 2014). CBLM whole-lobule ROIs were created using the SUIT atlas (Diedrichsen et al., 2009) and localized to the right hemisphere. In addition, we included two cortical seeds, one in the DLPFC and one in the M1, to investigate CBLM-frontal connectivity. These cortical seeds were chosen in parallel to CBLM whole-brain connectivity results from our past work using a sample of 590 females and males **Table A2**, where Crus I was associated with the DLPFC and Lobule V with M1 across sexes (Ballard et al., 2022a).

Finally, we extracted first-level ROI-to-ROI correlation results for Crus I-to-DLPFC connectivity and Lobule V-to-M1 connectivity in the form of z-scores (**Figure 1**). Importantly, the DLPFC is implicated in cognition (Smucny et al., 2022) and M1 is involved in motor function (Sanes and Donoghue, 2000), and both cortices demonstrate changes with age (Bhandari et al., 2016; MacPherson et al., 2002); thus, these cortical seeds also align with the functional properties of their respective CBLM regions.

**Figure 1.**
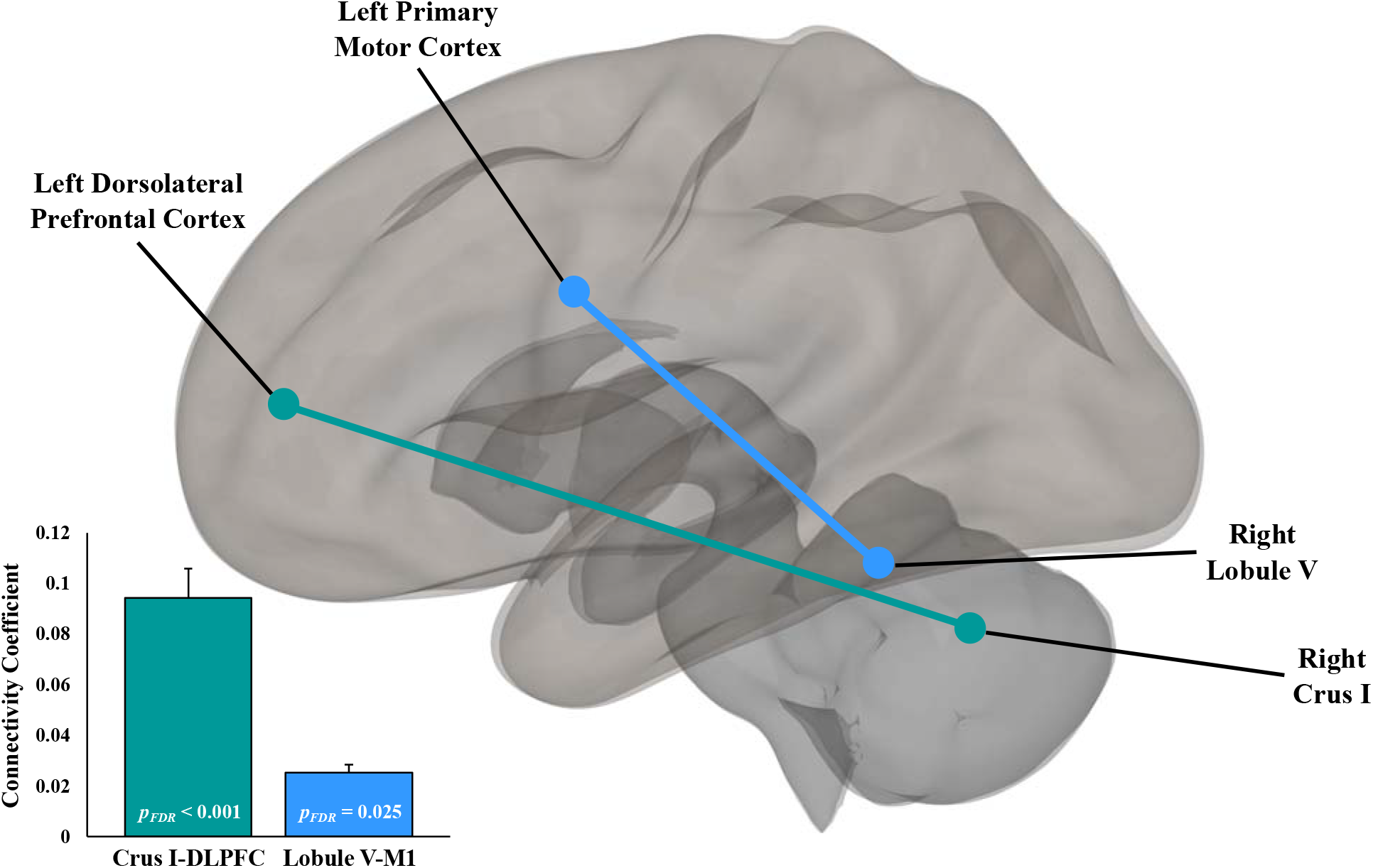
Cerebellar-Cortical Connections. Correlation coefficients and FDR-corrected p-values for ROI-to-ROI connectivity within each network in the current sample. For both networks, connectivity in the cerebellar and cortical node was significantly correlated. Error bars = standard error (SE).

### 2.8 Statistical Analyses

We first conducted a series of ANOVAs to investigate hormone differences between female reproductive stages. After including all hormones in a 3 × 3 ANOVA (reproductive stage by hormone type), we performed follow-up one-way ANOVAs for each hormone type. To test the influence of menopause, hormones, and sleep on the brain and behavior, we then performed two multivariate multiple linear regression models in R Studio (R Core Team, 2018). In Model 1, we used reproductive stage, 17*β*-estradiol, progesterone, testosterone, sleep quantity, sleep quality, and their interactions to predict Crus I- to-DLPFC connectivity and Lobule V-to-M1 connectivity. In Model 2, we used reproductive stage, 17*β*-estradiol, progesterone, testosterone, sleep quantity, sleep quality, Crus I-to-DLPFC connectivity, Lobule V-to-M1 connectivity, and their interactions to predict behavioral performance via the cognitive and motor composites. Statistical power was calculated *post-hoc* for each individual regression model using Cohen’s f^2^ (derived from R^2^_adj_) and an alpha level of 0.05 in G*Power (Cohen, 1988; Faul et al., 2007).

## 3. Results

### 3.1 Sex Hormone and Reproductive Stage Comparisons

We conducted a 3 × 3 between-subjects ANOVA with reproductive stage and hormone type as factors while hormone levels (pg/mL) were included as the outcome variable. This analysis revealed a significant effect of reproductive stage (*F*(2, 61) = 8.50, *p* < .001, *η*^2^ = 0.10) and hormone type (*F*(2, 122) = 57.24, *p* < .001, *η*^2^ = 0.35), and an interaction between reproductive stage and hormone type (*F*(4, 122) = 10.50, *p* < .001, *η*^2^ = 0.17).

We then conducted one-way ANOVAs with each hormone type to look for differences in individual hormone levels between stages. A significant difference in hormone levels between reproductive stages was present for both 17*β*-estradiol (*F*(2, 61) = 5.88, *p* = 0.005, *η*^2^ = 0.16) and progesterone (*F*(2, 61) = 12.13, *p* < .001, *η*^2^ = 0.28), but not for testosterone (*p* = 0.898). For 17*β*-estradiol and progesterone, hormone levels were lower with increasing stage from reproductive to postmenopausal, whereas testosterone levels remained stable across the menopausal transition (**Figure 2**).

**Figure 2.**
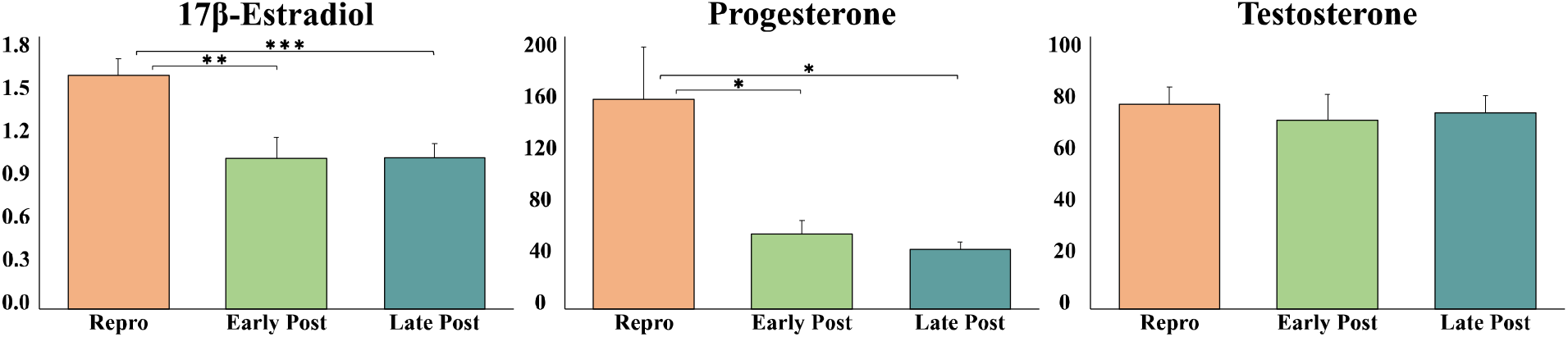
Hormone Levels (pg/mL) Across Groups. Repro: reproductive females; Early Post: early postmenopausal females; Late Post: late postmenopausal females. Error bars denote SE.

### 3.2 Crus I-to-DLPFC Connectivity

Results from the fitted model for Crus I-to-DLPFC connectivity (*F*(10, 37) = 5.48, *R*^2^_adj_ = 0.488, *p* < 0.001, *1-β* = 0.99) are presented in **Table B1** and **Figure 3**, and results from the original model are reported in **Table B2**. Crus I-to-DLPFC connectivity was significantly predicted by several variables (* = interaction): 17*β*-estradiol*sleep quantity (*β* = 0.110, *p* = 0.013), 17*β*-estradiol*sleep quality (*β* = -0.023, *p* = 0.030), progesterone*sleep quantity (*β* = -0.001, *p* = 0.022), and progesterone*sleep quality (*β* = 0.0001, *p* = 0.034).

**Figure 3.**
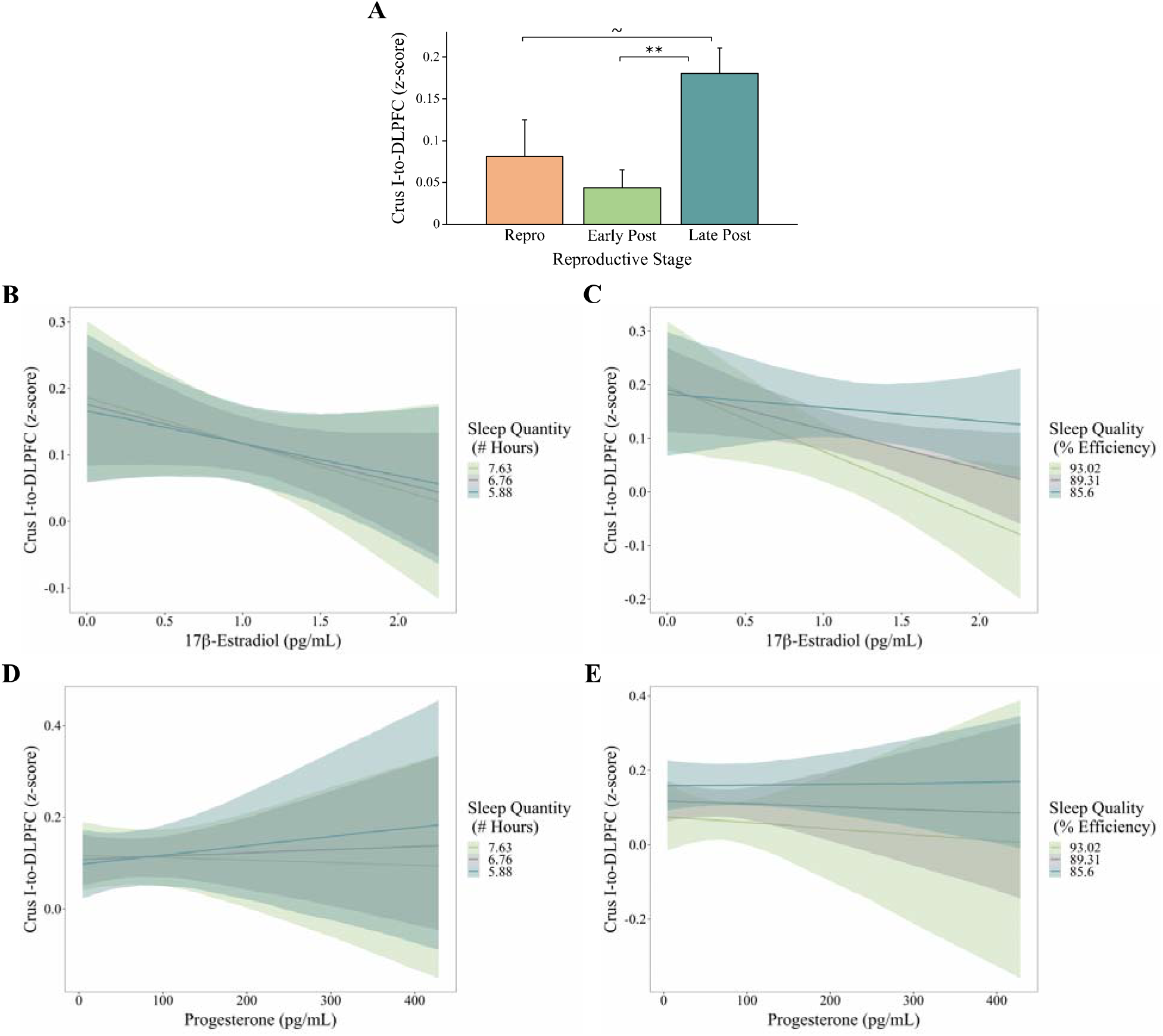
Significant Predictors for Crus I-to-DLPFC Connectivity. A) effect of reproductive stage; B) 17*β*-estradiol*sleep quantity interaction; C) 17*β*-estradiol*sleep quality interaction; D) progesterone*sleep quantity interaction; E) progesterone*sleep quality interaction. Repro: reproductive; Early Post: early postmenopause; Late Post: late postmenopause. Error bars represent SE. Grey: average slope and 95% confidence interval (CI); Green: projected slope and 95% CI for one SD above the mean; Blue: projected slope and 95% CI for one SD below the mean.

Further, a main effect of reproductive stage was present where late postmenopausal status predicted Crus I-to-DLPFC connectivity (*β* = 0.162, *p* < 0.001), relative to reproductive and early postmenopause. *Post-hoc* t-tests revealed a significant difference in connectivity between early and late postmenopausal females (*t*(33.63) = -3.71, *p* < 0.001), where late postmenopausal females displayed higher connectivity values. In addition, a trending difference between reproductive and late postmenopausal females in Crus I-to-DLPFC connectivity was also present (*t*(19.39) = -1.86, *p* = 0.078), with lower connectivity in the reproductive group. There were no significant differences however, between reproductive and early postmenopausal females (*p* = 0.460).

Compared to reproductive and early postmenopause groups, the late postmenopausal group was associated with higher Crus I-to-DLPFC connectivity (**Figure 3A**). Interestingly, higher 17*β*-estradiol and higher sleep quantity predict lower Crus I-to-DLPFC connectivity (**Figure 3B**). Relatedly, higher 17*β*-estradiol and higher sleep quality produce the lowest Crus I-to-DLPFC connectivity outcome as well, relative to low 17*β*-estradiol slopes which yield higher connectivity in this circuit (**Figure 3C**). Results for progesterone*sleep interactions are mixed; however, high progesterone and high sleep predict the lowest Crus I-to-DLPFC connectivity values, compared to other slopes (**Figure 3D** and **Figure 3E**).

### 3.3 Lobule V-to-M1 Connectivity

Results from the fitted model for Lobule V-to-M1 connectivity (*F*(9, 38) = 1.84, *R*^2^_adj_ = 0.139, *p* = 0.092, *1-β* = 0.53) are illustrated in **Table B3** and **Figure 4**. Results from the original model can be found in **Table B4**. Notably, the best fit model for Lobule V-to-M1 connectivity was not statistically significant and sufficient power (*1-β* ≥ 0.8) was not achieved. Model fit did not improve when removing non-significant variables in a stepwise fashion. However, a few variables within the best fit model significantly predicted Lobule V-to-M1 connectivity: 17*β*-estradiol (*β* = 0.399, *p* = 0.017) and 17*β*-estradiol*sleep quantity (*β* = -0.062, *p* = 0.018). Reproductive status did not impact the outcome in this model (*ps* > 0.169).

**Figure 4.**
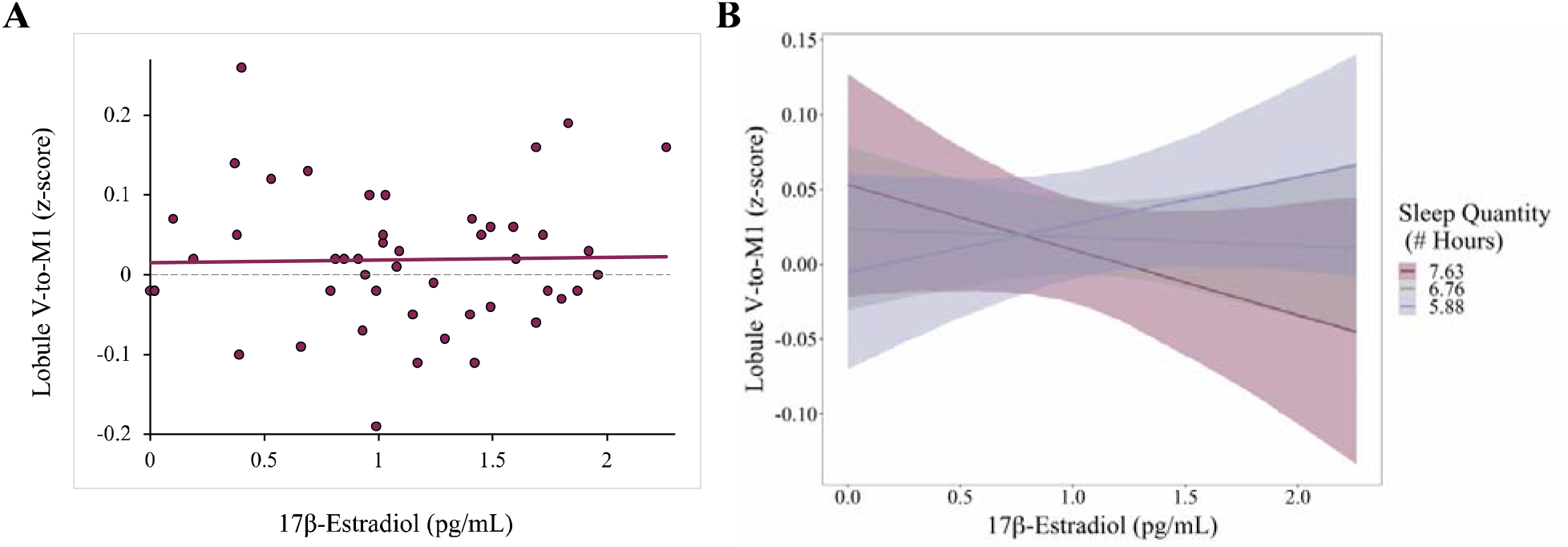
Significant Predictors for Lobule V-to-M1 Connectivity. A) effect of 17*β*-estradiol; B) 17*β*-estradiol*sleep quantity interaction. Grey: average slope and 95% CI; Magenta: projected slope and 95% CI for one SD above the mean; Purple: projected slope and 95% CI for one SD below the mean.

As 17*β*-estradiol increases, Lobule V-to-M1 connectivity also increases across females (**Figure 4A**). However, this relationship is modified by sleep quantity wherein those with lower sleep show higher Lobule V-to-M1 connectivity with higher 17*β*-estradiol. On the other hand, those with average sleep do not demonstrate much change in connectivity with increases in 17*β*-estradiol, while those with higher sleep show reductions in Lobule V-to-M1 connectivity in response to increases in 17*β*-estradiol (**Figure 4B**).

### 3.4 Cognitive Performance

Results from the fitted model for cognitive performance (*F*(18, 39) = 3.90, *R*^2^_adj_ = 0.478, *p* < 0.001, *1-β* = 0.99) are presented in **Table B5** and **Figure 5**, and results from the original model are provided in **Table B6**. For cognitive performance, as indexed by the average of standardized scores on three cognitive tasks, the following predictors were significant: testosterone (*β* = -0.010, *p* = 0.005), 17*β*-estradiol*sleep quantity (*β* = 0.518, *p* = 0.034), progesterone*sleep quantity (*β* = -0.004, *p* = 0.035), and sleep quantity*sleep quality (*β* = -0.063, *p* = 0.047). Further, an exploratory simple linear regression model revealed that 17*β*-estradiol significantly predicts cognitive behavior across females (*F*(1, 75) = 12.04, *R*^2^_adj_ = 0.127, *p* < 0.001, *β* = 0.436).

**Figure 5.**
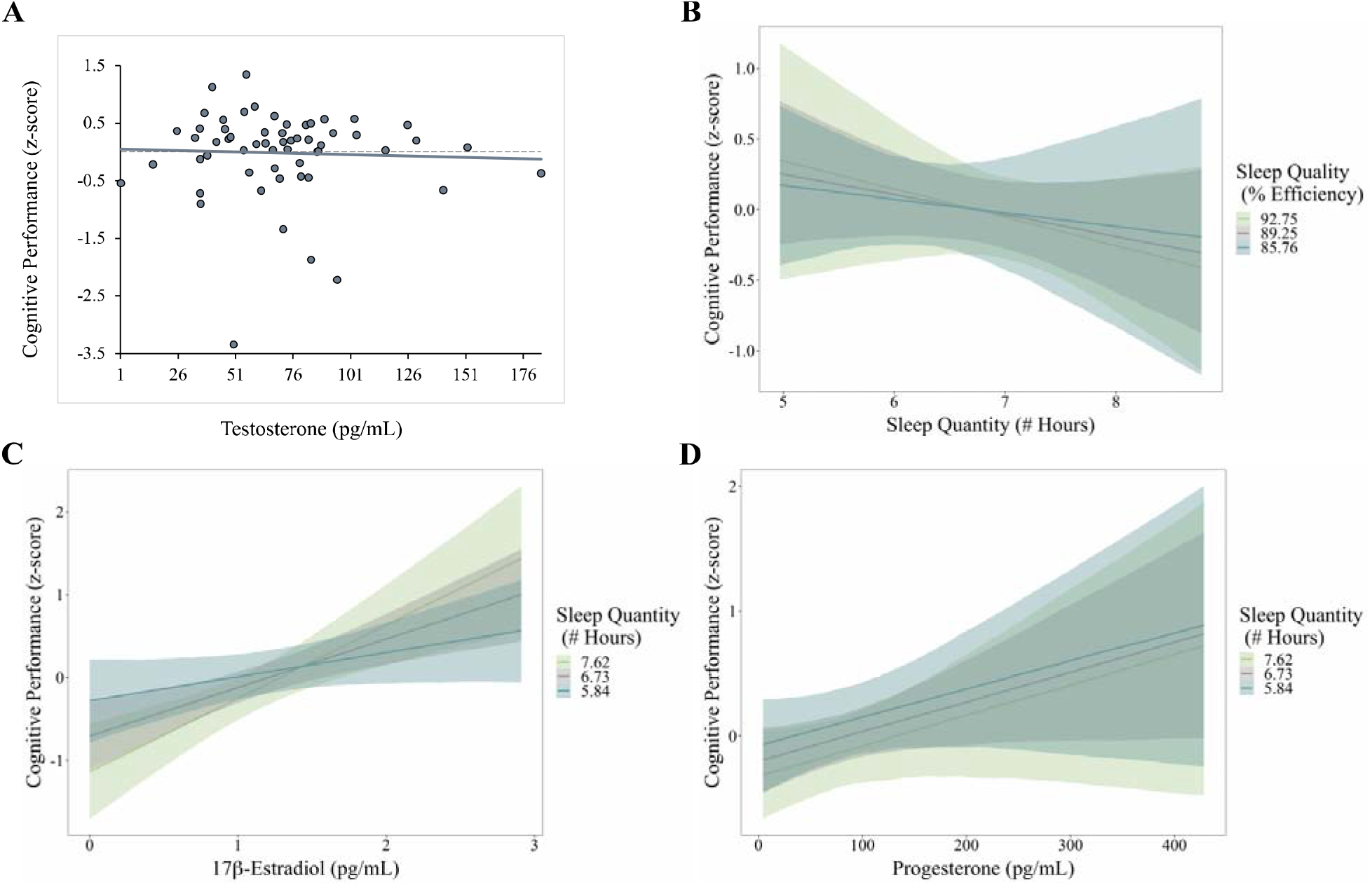
Significant Predictors for Cognitive Performance. A) effect of testosterone; B) sleep quantity*sleep quality interaction; C) 17*β*-estradiol*sleep quantity interaction; D) progesterone*sleep quantity interaction. Grey: mean slope and 95% CI; Green: projected slope and 95% CI for one SD above mean; Blue: projected slope and 95% CI for one SD below mean.

As testosterone increases, cognitive performance decreases across females (**Figure 5A**). Moreover, higher sleep quality, particularly in those with lower sleep quantity, produces the best cognitive outcome (**Figure 5B**). Further, the impact of hormones on cognition is also mediated by sleep quantity. Higher 17*β*-estradiol and higher sleep quantity produce the highest cognitive composite score (**Figure 5C**), though 17*β*-estradiol improves cognition in moderate and low sleepers as well. Paralleling these relationships, higher progesterone also produces better cognitive performance for all sleep quantity projections, whereas lower progesterone yields below-average cognitive scores (**Figure 5D**).

### 3.5 Motor Performance

Full results from the fitted model for motor performance (*F*(7, 65) = 4.16, *R*^2^_adj_ = 0.235, *p* < 0.001, *1-β* = 0.94) are depicted in **Table B7** and **Figure 6**. Results from the original model can be found in **Table B8**. All of the predictors incorporated into the best fit model for motor performance (average of z-scores on three motor tasks) were significant. Significant predictors include 17*β*-estradiol (*β* = 8.013, *p* = 0.006), testosterone (*β* = -0.008, *p* = 0.002), sleep quantity (*β* = 4.660, *p* = 0.037), sleep quality (*β* = 0.510, *p* = 0.005), 17*β*-estradiol*sleep quantity (*β* = 0.304, *p* = 0.026), 17*β*-estradiol*sleep quality (*β* = -0.107, *p* = 0.015), and sleep quantity*sleep quality (*β* = -0.056, *p* = 0.027).

**Figure 6.**
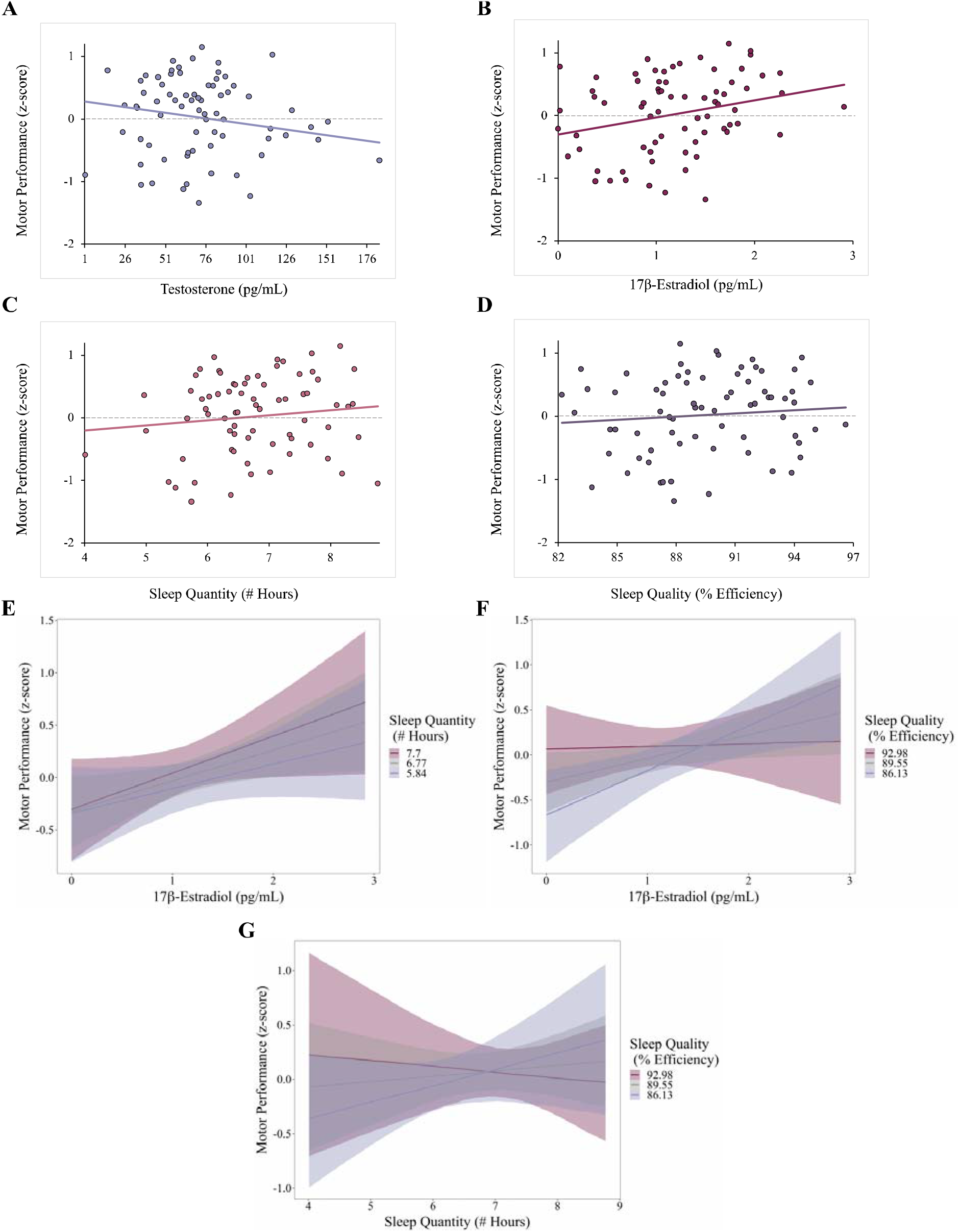
Significant Predictors for Motor Performance. A) effect of testosterone; B) effect of 17*β*-estradiol; C) effect of sleep quantity; D) effect of sleep quality; E) 17*β*-estradiol*sleep quantity interaction; F) 17*β*-estradiol*sleep quality interaction; G) sleep quantity*sleep quality interaction. Grey: mean slope and 95% CI; Magenta: projected slope and 95% CI for one SD above mean; Purple: projected slope and 95% CI for one SD below mean.

Similar to the cognitive composite, lower motor performance scores were associated with higher testosterone in females (**Figure 6A**). On the other hand, higher motor scores are predicted by higher 17*β*-estradiol (**Figure 6B**), higher sleep quantity (**Figure 6C**), and higher sleep quality (**Figure 6D**). With higher sleep quantity, higher 17*β*-estradiol predicts the best motor outcome, and the benefit of 17*β*-estradiol on motor performance also emerges for moderate and low sleep quantity groups (**Figure 6E**). However, for those with moderate and lower sleep quality, increases in 17*β*-estradiol improve motor outcomes (**Figure 6F**). Thus, it may be the case that 17*β*-estradiol helps buffer against the impact of low sleep quality on motor behavior. Finally, sleep quantity and quality interact to predict motor performance as higher sleep quality is associated with higher scores in those with lower sleep quantity, mirroring the same suspected sleep trade-off observed with cognitive outcomes (**Figure 6G**).

## 4. Discussion

### 4.1 Summary of Findings

We used salivary hormone levels and actigraphy-derived sleep assessments to predict cognitive/motor behavior and CBLM-frontal connectivity in healthy MA and OA females. First considering our results with CBLM-frontal connectivity, we found somewhat dissociable effects between cognitive (Crus I-to-DLPFC) and motor (Lobule V-to-M1) networks. For both networks, 17*β*-estradiol and sleep quantity interacted to predict connectivity, though the response of each network to these factors was unique. Moreover, significant effects of progesterone-sleep interactions were observed with the Crus I-to-DLPFC network, though impacts of 17*β*-estradiol and progesterone partially diverge. That is, 17*β*-estradiol and sleep maintain a positive association (increase together), while progesterone-sleep relationships are bi-directional. Finally, differences in connectivity between reproductive stages emerged for the Crus I-to-DLPFC network, while Lobule V-to-M1 connectivity did not present group differences.

Both aspects of behavior were also impacted by hormone-sleep interactions, irrespective of menopausal status. We observed a small inhibitory effect of testosterone on behavioral performance, while interactions between 17*β*-estradiol and sleep quantity/quality were related to better cognitive and motor outcomes. Interestingly, sleep quantity and quality, when taken together, do not have linear effects on cognition. It seems that a specific combination of the number of hours slept and time slept versus time spent in bed is necessary for optimal behavior. Unique to motor function, we also observed several main effects of hormone and sleep variables, paralleling the observed interactions between these factors.

Collectively, our results suggest that CBLM-frontal networks are dynamically impacted by menopausal status and hormone-sleep interactions, however a few consistencies also arise. On the contrary, distinct domains of behavior demonstrate parallel responses to sex hormones and sleep patterns, though some slight differences were present there as well.

### 4.2 Cerebellar-Frontal Networks are Susceptible to the Effects of Hormone-Sleep Interactions

Though we anticipated reduced CBLM-frontal connectivity for postmenopausal females relative to reproductive, our findings of higher connectivity in the Crus I-to-DLPFC network are consistent with some of the relevant work in this area. In our own prior work, we found that nearly 70% of differences between premenopausal and postmenopausal females were due to lower CBLM connectivity in the postmenopausal group. However, we also observed a few instances where connectivity was greater in this group, parallel to what we report here. For example, we found greater connectivity between cognitive constituents of the CBLM (Crus I/II) and occipital/parietal cortices, as well as between motor CBLM lobules (Lobules V-VI) and the prefrontal cortex, in postmenopausal females compared to reproductive (Ballard et al., 2022a). Fitzgerald et al. (2020) reports both positive and negative associations of estradiol and progesterone with CBLM connectivity, suggesting that CBLM-cortical networks are dynamically reactive to sex hormones. Oscillations in network organization and efficiency are consistently observed with hormone fluctuations across the menstrual cycle (Pritschet et al., 2020); thus, the hormonal profile of menopause may, in turn, produce wide-ranging effects on brain networks, as observed here.

Higher connectivity, as seen in the Crus I-to-DLPFC network in our postmenopausal sample, is associated with age-related events, such as the onset of AD. Phases of hypoconnectivity/hyperconnectivity are commonly reported with the progression of AD (Yao et al., 2021), and these changes are observed with CBLM-cortical networks as well (Tang et al., 2021). As such, greater network connectivity may represent a global response to neurological dysfunction, which may relate to the rapid destabilization of the hormonal equilibrium with the transition to menopause. Between-network hyperconnectivity may also reflect a compensatory mechanism to reduced efficiency within networks, contributing to the dedifferentiation of large-scale functional organization in older age (Cabeza et al., 2018). In previous work, we found that reductions in functional network segregation with age are sex-specific in nature; relative to males, females drive negative age-segregation associations for the dorsal attention and default mode networks, whereas males show a unique relationship with the somatosensory hand network (Ballard et al., 2022b). Pertinent to the current findings, the dorsal attention and default mode networks are comprised of prefrontal areas that correspond to that of our Crus I-to-DLPFC network, which also displayed female-specific traits. Given variable changes in the functional architecture of the brain with age, it may be the case that increased CBLM-cortical connectivity with endocrine aging is an attempt to restructure the brain after hormones have reached a stabilized low, though this explanation is purely speculative.

Interactions between sex hormones and sleep patterns also impacted our connectivity outcomes. Considering the literature supporting a relationship between hormones and sleep (Brown and Gervais, 2020), this finding was largely anticipated. Several potential mechanisms for the neural linkage between hormones and sleep have been proposed in both animal and human studies, including hormonal interactions with circadian rhythms and the memory-promoting benefits of sleep (Brown and Gervais, 2020). Though more detailed investigations are needed, our results build upon an established relationship between these age-related factors by assessing their collective impact on CBLM-frontal networks and behavior.

### 4.3 Interactions Between Sex Hormones and Sleep Patterns Impact Behavior

While estrogen has a reputation for enhancing cognition (Ghidoni et al., 2006), findings with progesterone are mixed. Some trials record improvements in cognition and sleep with progesterone therapy in postmenopausal females (Berent-Spillson et al., 2015), whereas others illustrate null or negative effects (Henderson, 2018). Our results follow suit, illustrating similar yet alternating impacts of estrogen and progesterone on female behavior. Surprisingly, we observed negative effects of testosterone on both cognitive and motor performance across females and irrespective of menopausal status. Similar to progesterone, the impact of testosterone on behavior in females is inconsistent, and work in this area is limited. However, akin to what emerged in our sample, van Honk et al. (2011) notes impaired cognition in young females with testosterone administration. Together, our findings substantiate the claim that fluctuating hormones enact a wide range of varying effects on the female brain and behavior in later life.

Though more sleep is commonly linked to better behavioral performance, our findings indicate that the relationship between sleep quantity and quality may not be linear. With both behavioral domains, we found a trade-off between sleep quantity and quality where an optimal combination seems to produce the best behavioral outcomes. We see a shift in the effect of sleep quality on behavior at about 7 hours of sleep quantity, where higher quality past this point does not predict better behavior but lower quality groups do show a benefit from additional sleep hours. In fact, some work describes a negative effect of excessive sleep; a study by Wild et al. (2018) with 10,000+ participants shows that those who slept ± 7-8 hours, on average, exhibited worsened cognitive performance across multiple tests. This suggests that there is a somewhat limited range of optimal sleep with respect to the impact on cognition. Ferrie et al. (2011) supports this notion, illustrating that an increase in sleep between baseline and follow-up to an amount that is in excess of the average 7-8 hours/night is accompanied by broad declines in cognition. Thus, though initially unanticipated, our results broadly parallel other work suggesting that the benefits of sleep are not completely linear but rather an optimal level exists.

Finally, we did not observe a predictive effect of CBLM-frontal connectivity on behavior in our analyses. While Crus I-to-DLPFC and Lobule V-to-M1 connectivity were included in the original models, these variables were removed upon adjustment. Though Crus I and the DLPFC have purported roles in cognition and Lobule V/M1 in motor function (O’Reilly et al., 2010), these specific networks may not globally subserve cognitive and motor behavior in MA and OA females, as these outcomes were represented by a composite average of scores on three unique tasks per domain in our study. Thus, certain components of these behavioral composites may be driven by different networks. A broader exploration of CBLM-frontal connectivity may provide better insight on the contribution of these networks, and others, to behavior in adulthood.

### 4.4 Experimental Limitations

Several limitations are worth noting. First, we were not able to evaluate the perimenopausal stage in our analyses as only 3 females were determined to be in this stage. Further, our reproductive stage classifications were based off of self-report data over the course of 6 months, whereas the STRAW+10 criteria recommend categorizing females using 10 cycles-worth of data. We also did not collect data regarding day of cycle for reproductive females, though different phases of the menstrual cycle have varying concentrations of sex hormones, which may have contributed to the lack of differences relative to menopausal status with some outcome variables. Salivary samples were collected on the day of behavioral testing, which occurred a few weeks prior to the MRI session; thus, reproductive females could have potentially entered a different phase of their cycle by the time of resting-state data collection, and this could have additionally impacted group differences.

Second, as our experimental procedures were split between two separate sessions, the unanticipated restrictions on data collection in response to the Covid-19 pandemic caused the time between sessions to vary more than intended across subjects (average time between sessions = 38 days). We also did not exclude left-handed (n = 0) or ambidextrous (n = 2) subjects from this study; thus, potential differences in brain organization due to handedness may have impacted results, though this only affected 3% of our sample. However, our CBLM-cortical networks traverse both hemispheres, helping to partially offset variations in lateralization of function. This, in turn, results in a sample that is more representative of the population.

With regards to sleep quantity, we did not screen participants for possible sleep disorders that are accompanied by increased movement during sleep (e.g., REM sleep behavior disorder or sleep apnea), which may be relevant to the detection of sleep periods and classification of sleep fragmentation in our sample. Moreover, the actigraphy-based sleep quality variable is only a measure of the proportion of time slept relative to time spent in bed, which is sufficient for measuring sleep efficiency but does not provide detailed information on sleep stages, (e.g., non-REM versus REM), which is an important distinction for establishing the quality of sleep. Though actigraphy recordings are well-correlated with polysomnographic variables (Quante et al., 2018), the actigraphy method still lacks some of the nuance that is achieved with polysomnography, and this may limit the interpretability of our sleep results.

### 4.5 Conclusions

The current work represents a novel interrogation of sex hormones and sleep patterns, exploring their role in brain and behavioral outcomes over the course of female adulthood. Using accessible, non-invasive measures, we probed the interaction between hormones and sleep to determine the influence of these age-linked factors on CBLM-frontal networks and cognitive/motor behavior. This investigation provides new insight into the neural mechanisms and biological variables that may contribute to the divergent aging trajectories observed between females and males. Such information may be useful in developing strategies and targeted interventions for addressing functional declines in later life, particularly those that disproportionately impact older females.

## Funding

This work was supported by the National Institute on Aging (grant number R01AG065010).

## Notes

### Competing Interest Statement

The authors have declared no competing interest.

## References

Alzheimer’s Association, 2021. 2021 Alzheimer’s disease facts and figures. Alzheimers Dement. 17, 327–406. doi:10.1002/alz.12328

Anstey, K.J., Peters, R., Mortby, M.E., Kiely, K.M., Eramudugolla, R., Cherbuin, N., Huque, M.H., Dixon, R.A., 2021. Association of sex differences in dementia risk factors with sex differences in memory decline in a population-based cohort spanning 20-76 years. Sci. Rep. 11, 7710. doi:10.1038/s41598-021-86397-7

Ballard, H.K., Jackson, T.B., Hicks, T.H., Bernard, J.A., 2022a. The association of reproductive stage with lobular cerebellar network connectivity across female adulthood. Neurobiol. Aging 117, 139–150. doi:10.1016/j.neurobiolaging.2022.05.014

Ballard, H.K., Jackson, T.B., Miller, A.C., Hicks, T.H., Bernard, J.A., 2022b. Age-related differences in functional network segregation in the context of sex and reproductive stage. BioRxiv. doi:10.1101/2022.03.28.486067

Benson, N., Hulac, D.M., Kranzler, J.H., 2010. Independent examination of the Wechsler Adult Intelligence Scale-Fourth Edition (WAIS-IV): what does the WAIS-IV measure? Psychol Assess 22, 121–130. doi:10.1037/a0017767

Berent-Spillson, A., Briceno, E., Pinsky, A., Simmen, A., Persad, C.C., Zubieta, J.-K., Smith, Y.R., 2015. Distinct cognitive effects of estrogen and progesterone in menopausal women. Psychoneuroendocrinology 59, 25–36. doi:10.1016/j.psyneuen.2015.04.020

Bernard, J.A., Seidler, R.D., 2014. Moving forward: age effects on the cerebellum underlie cognitive and motor declines. Neurosci. Biobehav. Rev. 42, 193–207. doi:10.1016/j.neubiorev.2014.02.011

Bhandari, A., Radhu, N., Farzan, F., Mulsant, B.H., Rajji, T.K., Daskalakis, Z.J., Blumberger, D.M., 2016. A meta-analysis of the effects of aging on motor cortex neurophysiology assessed by transcranial magnetic stimulation. Clin. Neurophysiol. 127, 2834–2845. doi:10.1016/j.clinph.2016.05.363

Brown, A.M.C., Gervais, N.J., 2020. Role of ovarian hormones in the modulation of sleep in females across the adult lifespan. Endocrinology 161. doi:10.1210/endocr/bqaa128

Cabeza, R., Albert, M., Belleville, S., Craik, F.I.M., Duarte, A., Grady, C.L., Lindenberger, U., Nyberg, L., Park, D.C., Reuter-Lorenz, P.A., Rugg, M.D., Steffener, J., Rajah, M.N., 2018. Maintenance, reserve and compensation: the cognitive neuroscience of healthy ageing. Nat. Rev. Neurosci. 19, 701–710. doi:10.1038/s41583-018-0068-2

Canto, C.B., Onuki, Y., Bruinsma, B., van der Werf, Y.D., De Zeeuw, C.I., 2017. The Sleeping Cerebellum. Trends Neurosci. 40, 309–323. doi:10.1016/j.tins.2017.03.001

Cohen, J., 1988. Statistical power analysis for the behavioral sciences, 2nd ed. Lawrence Erlbaum Associates, New Jersey, NJ. doi:10.4324/9780203771587

Diedrichsen, J., Balsters, J.H., Flavell, J., Cussans, E., Ramnani, N., 2009. A probabilistic MR atlas of the human cerebellum. Neuroimage 46, 39–46. doi:10.1016/j.neuroimage.2009.01.045

Faul, F., Erdfelder, E., Lang, A.-G., Buchner, A., 2007. G*Power 3: A flexible statistical power analysis program for the social, behavioral, and biomedical sciences. Behav. Res. Methods 39, 175–191. doi:10.3758/BF03193146

Ferrie, J.E., Shipley, M.J., Akbaraly, T.N., Marmot, M.G., Kivimäki, M., Singh-Manoux, A., 2011. Change in sleep duration and cognitive function: findings from the Whitehall II Study. Sleep 34, 565–573. doi:10.1093/sleep/34.5.565

Fitzgerald, M., Pritschet, L., Santander, T., Grafton, S.T., Jacobs, E.G., 2020. Cerebellar network organization across the human menstrual cycle. Sci. Rep. 10, 20732. doi:10.1038/s41598-020-77779-4

Flegal, K.E., Lustig, C., 2016. You can go your own way: effectiveness of participant-driven versus experimenter-driven processing strategies in memory training and transfer. Neuropsychol Dev Cogn B Aging Neuropsychol Cogn 23, 389–417. doi:10.1080/13825585.2015.1108386

Ghidoni, R., Boccardi, M., Benussi, L., Testa, C., Villa, A., Pievani, M., Gigola, L., Sabattoli, F., Barbiero, L., Frisoni, G.B., Binetti, G., 2006. Effects of estrogens on cognition and brain morphology: involvement of the cerebellum. Maturitas 54, 222–228. doi:10.1016/j.maturitas.2005.11.002

Gip, P., Hagiwara, G., Ruby, N.F., Heller, H.C., 2002. Sleep deprivation decreases glycogen in the cerebellum but not in the cortex of young rats. Am. J. Physiol. Regul. Integr. Comp. Physiol. 283, R54–9. doi:10.1152/ajpregu.00735.2001

Hajali, V., Andersen, M.L., Negah, S.S., Sheibani, V., 2019. Sex differences in sleep and sleep loss-induced cognitive deficits: The influence of gonadal hormones. Horm. Behav. 108, 50–61. doi:10.1016/j.yhbeh.2018.12.013

Hara, Y., Waters, E.M., McEwen, B.S., Morrison, J.H., 2015. Estrogen effects on cognitive and synaptic health over the lifecourse. Physiol. Rev. 95, 785–807. doi:10.1152/physrev.00036.2014

Harlow, S.D., Gass, M., Hall, J.E., Lobo, R., Maki, P., Rebar, R.W., Sherman, S., Sluss, P.M., de Villiers, T.J., STRAW + 10 Collaborative Group, 2012. Executive summary of the Stages of Reproductive Aging Workshop + 10: addressing the unfinished agenda of staging reproductive aging. J. Clin. Endocrinol. Metab. 97, 1159–1168. doi:10.1210/jc.2011-3362

Harms, M.P., Somerville, L.H., Ances, B.M., Andersson, J., Barch, D.M., Bastiani, M., Bookheimer, S.Y., Brown, T.B., Buckner, R.L., Burgess, G.C., Coalson, T.S., Chappell, M.A., Dapretto, M., Douaud, G., Fischl, B., Glasser, M.F., Greve, D.N., Hodge, C., Jamison, K.W., Jbabdi, S., Kandala, S., Li, X., Mair, R.W., Mangia, S., Marcus, D., Mascali, D., Moeller, S., Nichols, T.E., Robinson, E.C., Salat, D.H., Smith, S.M., Sotiropoulos, S.N., Terpstra, M., Thomas, K.M., Tisdall, M.D., Ugurbil, K., van der Kouwe, A., Woods, R.P., Zöllei, L., Van Essen, D.C., Yacoub, E., 2018. Extending the Human Connectome Project across ages: Imaging protocols for the Lifespan Development and Aging projects. Neuroimage 183, 972–984. doi:10.1016/j.neuroimage.2018.09.060

Hedges, V.L., Chen, G., Yu, L., Krentzel, A.A., Starrett, J.R., Zhu, J.-N., Suntharalingam, P., Remage-Healey, L., Wang, J.-J., Ebner, T.J., Mermelstein, P.G., 2018. Local Estrogen Synthesis Regulates Parallel Fiber-Purkinje Cell Neurotransmission Within the Cerebellar Cortex. Endocrinology 159, 1328–1338. doi:10.1210/en.2018-00039

Henderson, V.W., 2018. Progesterone and human cognition. Climacteric 21, 333–340. doi:10.1080/13697137.2018.1476484

Kwak, Y., Müller, M.L.T.M., Bohnen, N.I., Dayalu, P., Seidler, R.D., 2012. l-DOPA changes ventral striatum recruitment during motor sequence learning in Parkinson’s disease. Behav. Brain Res. 230, 116–124. doi:10.1016/j.bbr.2012.02.006

LeBlanc, E.S., Janowsky, J., Chan, B.K., Nelson, H.D., 2001. Hormone replacement therapy and cognition: systematic review and meta-analysis. JAMA 285, 1489–1499. doi:10.1001/jama.285.11.1489

Levine, D.A., Gross, A.L., Briceño, E.M., Tilton, N., Giordani, B.J., Sussman, J.B., Hayward, R.A., Burke, J.F., Hingtgen, S., Elkind, M.S.V., Manly, J.J., Gottesman, R.F., Gaskin, D.J., Sidney, S., Sacco, R.L., Tom, S.E., Wright, C.B., Yaffe, K., Galecki, A.T., 2021. Sex differences in cognitive decline among US adults. JAMA Netw. Open 4, e210169. doi:10.1001/jamanetworkopen.2021.0169

Li, D.X., Romans, S., De Souza, M.J., Murray, B., Einstein, G., 2015. Actigraphic and self-reported sleep quality in women: associations with ovarian hormones and mood. Sleep Med. 16, 1217–1224. doi:10.1016/j.sleep.2015.06.009

Li, L., Nakamura, T., Hayano, J., Yamamoto, Y., 2021. Age and gender differences in objective sleep properties using large-scale body acceleration data in a Japanese population. Sci. Rep. 11, 9970. doi:10.1038/s41598-021-89341-x

MacPherson, S.E., Phillips, L.H., Della Sala, S., 2002. Age, executive function and social decision making: A dorsolateral prefrontal theory of cognitive aging. Psychol. Aging 17, 598–609. doi:10.1037/0882-7974.17.4.598

Markowska, A.L., 1999. Sex dimorphisms in the rate of age-related decline in spatial memory: relevance to alterations in the estrous cycle. J. Neurosci. 19, 8122–8133.

Oliveira, L.F., Simpson, D.M., Nadal, J., 1996. Calculation of area of stabilometric signals using principal component analysis. Physiol Meas 17, 305–312. doi:10.1088/0967-3334/17/4/008

O’Reilly, J.X., Beckmann, C.F., Tomassini, V., Ramnani, N., Johansen-Berg, H., 2010. Distinct and overlapping functional zones in the cerebellum defined by resting state functional connectivity. Cereb. Cortex 20, 953–965. doi:10.1093/cercor/bhp157

Pritschet, L., Santander, T., Taylor, C.M., Layher, E., Yu, S., Miller, M.B., Grafton, S.T., Jacobs, E.G., 2020. Functional reorganization of brain networks across the human menstrual cycle. Neuroimage 220, 117091. doi:10.1016/j.neuroimage.2020.117091

Quante, M., Kaplan, E.R., Cailler, M., Rueschman, M., Wang, R., Weng, J., Taveras, E.M., Redline, S., 2018. Actigraphy-based sleep estimation in adolescents and adults: a comparison with polysomnography using two scoring algorithms. Nat. Sci. Sleep 10, 13–20. doi:10.2147/NSS.S151085

R Core Team, 2018. R Studio.

Rehbein, E., Hornung, J., Sundström Poromaa, I., Derntl, B., 2021. Shaping of the female human brain by sex hormones: A review. Neuroendocrinology 111, 183–206. doi:10.1159/000507083

Sabia, S., Fayosse, A., Dumurgier, J., van Hees, V.T., Paquet, C., Sommerlad, A., Kivimäki, M., Dugravot, A., Singh-Manoux, A., 2021. Association of sleep duration in middle and old age with incidence of dementia. Nat. Commun. 12, 2289. doi:10.1038/s41467-021-22354-2

Salimetrics, L., 2006. High sensitivity salivary estradiol enzyme immunoassay kit: performance characteristics. State College (PA), unpublished author.

Sanes, J.N., Donoghue, J.P., 2000. Plasticity and primary motor cortex. Annu. Rev. Neurosci. 23, 393–415. doi:10.1146/annurev.neuro.23.1.393

Smucny, J., Dienel, S.J., Lewis, D.A., Carter, C.S., 2022. Mechanisms underlying dorsolateral prefrontal cortex contributions to cognitive dysfunction in schizophrenia. Neuropsychopharmacology 47, 292–308. doi:10.1038/s41386-021-01089-0

Stroop, J.R., 1935. Studies of interference in serial verbal reactions. J Exp Psychol 18, 643–662. doi:10.1037/h0054651

Tang, F., Zhu, D., Ma, W., Yao, Q., Li, Q., Shi, J., 2021. Differences Changes in Cerebellar Functional Connectivity Between Mild Cognitive Impairment and Alzheimer’s Disease: A Seed-Based Approach. Front. Neurol. 12, 645171. doi:10.3389/fneur.2021.645171

Thomas, M., Sing, H., Belenky, G., Holcomb, H., Mayberg, H., Dannals, R., Wagner, H., Thorne, D., Popp, K., Rowland, L., Welsh, A., Balwinski, S., Redmond, D., 2000. Neural basis of alertness and cognitive performance impairments during sleepiness. I. Effects of 24 h of sleep deprivation on waking human regional brain activity. J Sleep Res 9, 335–352. doi:10.1046/j.1365-2869.2000.00225.x

Tiffin, J., Asher, E.J., 1948. The Purdue pegboard; norms and studies of reliability and validity. J. Appl. Psychol. 32, 234–247. doi:10.1037/h0061266

van Honk, J., Schutter, D.J., Bos, P.A., Kruijt, A.-W., Lentjes, E.G., Baron-Cohen, S., 2011. Testosterone administration impairs cognitive empathy in women depending on second-to-fourth digit ratio. Proc. Natl. Acad. Sci. USA 108, 3448–3452. doi:10.1073/pnas.1011891108

Whitfield-Gabrieli, S., Nieto-Castanon, A., 2012. Conn: a functional connectivity toolbox for correlated and anticorrelated brain networks. Brain Connect. 2, 125–141. doi:10.1089/brain.2012.0073

Wild, C.J., Nichols, E.S., Battista, M.E., Stojanoski, B., Owen, A.M., 2018. Dissociable effects of self-reported daily sleep duration on high-level cognitive abilities. Sleep 41. doi:10.1093/sleep/zsy182

Yao, W., Chen, H., Luo, C., Sheng, X., Zhao, H., Xu, Y., Bai, F., Alzheimer’s Disease Neuroimaging Initiative, 2021. Hyperconnectivity of Self-Referential Network as a Predictive Biomarker of the Progression of Alzheimer’s Disease. J. Alzheimers Dis. 80, 577–590. doi:10.3233/JAD-201376

