## Appendix A for "Hormone-sleep interactions predict cerebellar connectivity and behavior in aging females"

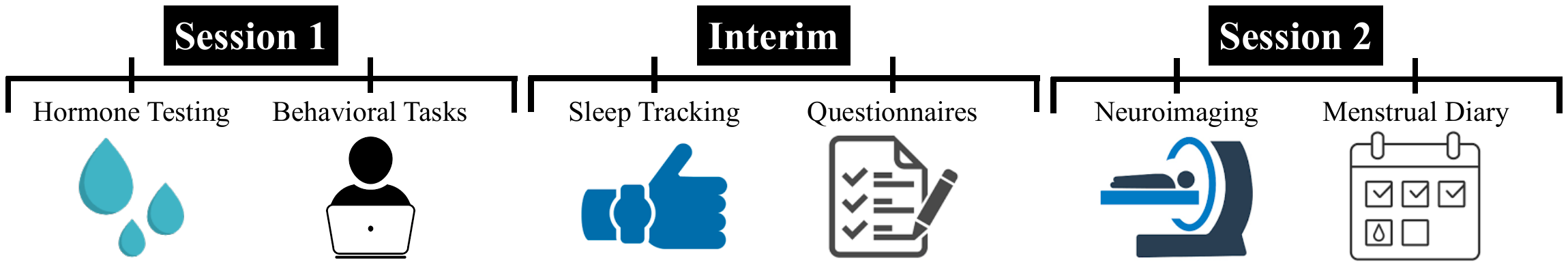

Figure A1. *Schematic Overview of Experimental Procedures.*

*A1. Hormone Testing*

To avoid exogenous influences on hormone levels, participants were asked to refrain from consuming alcohol 24 hours prior and eating or drinking 3 hours prior to their first study session. Participants were also screened for oral disease or injury, use of nicotine or caffeine, and prescription medications that may impact the saliva pH and compromise samples. Participants were asked to rinse their mouth with water 10 minutes prior to providing a saliva sample to clear out any residue.

One late postmenopausal female, age 60, had an average 17$\beta$-estradiol reading that was 5 standard deviations (SDs) above their respective stage group mean and 6 SDs above the overall mean across females. This participant was excluded from the final analyses in the interest of caution. We did not detect any additional hormonal outliers across hormone types in terms of $\pm$ 3 SDs from the overall or group means.

*A2. Reproductive Staging*

Though the STRAW+10 criteria recommends classifying females using data across 10 consecutive cycles, we only recorded information for 6 cycles to increase ease for participants. The consistency and completeness of responses on the menstrual tracking diary varied; thus, we did not gain enough information to accurately categorize 11 participants (ages 39-68).

Several participants reported a history of hysterectomy but were still included in their respective stage group (n = 4, 1 early postmenopausal, 3 late postmenopausal). These individuals underwent their procedures at least 8 years prior to participating in our study, with the majority being 30+ years earlier. We kept these participants in our analyses as hormone levels have likely reached a stabilized low, putting them in a comparable state to their naturally menopausal counterparts.

Table A1. *Characteristics for Female Stage Groups.*

| *Stage* | *Sample Size* | *Mean Age* | *Age Range* |
| --- | --- | --- | --- |
| Reproductive | 15 | 41.00 $\pm$ 4.57 | 35-49 |
| Early Postmenopause | 18 | 56.17 $\pm$ 4.15 | 47-62 |
| Late Postmenopause | 35 | 68.94 $\pm$ 8.22 | 52-86 |

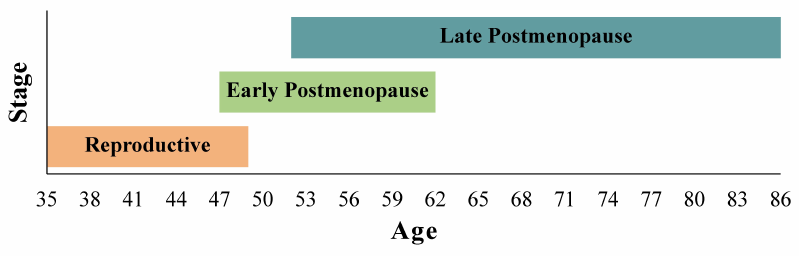

Figure A2. *Group Age Range Distributions.*

*A3. Cognitive Tasks*

During the Stroop Task, participants were presented with a color word (i.e., red, blue, or green) in colored ink. Participants were then asked to respond to the ink color of each word, rather than the word itself. For example, if a participant was presented with the word “blue” in green ink, they were asked to respond “green” to the trial using designated keys on a computer keyboard (i.e., “g” for green ink, “r” for red ink, or “b” for blue ink). Following a brief practice period that provided feedback on their responses, participants completed a test phase with 10 blocks. Each block contained 18 congruent trials where the ink color and color word were the same (e.g., the word “blue” presented in blue ink) and 18 incongruent trials where the color word was presented in an alternative ink color (e.g., the word “blue” presented in red ink). Thus, a total of 360 trials, 50% congruent and 50% incongruent, were included in the task. Given that one participant did not fully complete the Stroop Task and another missed over 90% of trials on the Stroop Task, the composite score for these two participants was only averaged between the Shopping List Memory Task and Digit-Symbol Substitution Task.

For the Shopping List Memory Task, participants viewed 15 items from a grocery list (e.g., milk, bread, eggs – see Appendix A) that were presented one at a time for 3 seconds each during an encoding phase. After a 15-minute delay, participants then completed the retrieval phase. The retrieval phase included 15 lures, or items similar but not exact to those presented in the encoding phase, and 15 targets (exact same items from the encoding phase). Participants responded to each item as either “old” (target) or “new” (lure), relative to the encoding phase, using specific keys on the computer keyboard.

During the Digit-Symbol Substitution Task, participants were provided with a paper worksheet that contained several rows of digits with empty boxes below and a digit-symbol key. Each digit was assigned to a corresponding symbol, and participants were instructed to copy the appropriate symbol for each digit in the empty box below. Participants were given 2 minutes to complete as many items as possible, in order from left to right.

*A4. Motor Tasks*

For the Postural Sway Task, we used an Advanced Mechanical Technology Incorporated (AMTI) Accusway force platform (Watertown, MA), which quantifies postural sway by recording deviations from the center of pressure (COP). Participants completed four trials, each lasting 2 minutes, where they were asked to stand as still as possible in the center of the plate for four different positions that varied visual input and foot placement: eyes open/feet shoulder apart, eyes closed/feet apart, eyes open/feet together, and eyes closed/feet together. For each position, participants were also asked to cross their arms over their chest and fixate on a central cross placed on the opposite wall. One subject did not have data for 1/4 positions (eyes open/feet together); thus, their average score was calculated using only three positions. In addition, one subject was excluded entirely from calculations for this task as their COP area was 9 SDs above the group mean, suggesting that a technical error may have occurred.

The Purdue Pegboard Task used a pegboard with two parallel columns of 25 holes. At the top of the pegboard were pins/pegs in the leftmost and rightmost bins and small screws or washers in the middle bins. Participants first completed a trial in which they were instructed to place the pegs from the left bin into the left column using their left hand. Next, participants were asked to use their right hand to place the pegs form the right bin into the right column. Third, participants were asked to use both hands to place pegs from both the left and right bins into a row of holes simultaneously. For each of these first three trials, participants were given 30 seconds to complete as many placements as possible (from top to bottom row). Lastly, participants completed an assembly trial, using all four items in the following order: peg, washer, screw, washer. During this trial, participants were instructed to alternate hands and were given 60 seconds to complete as many assemblies as possible. Finally, all four trials (i.e., left hand/left column, right hand/right column, both hands/both columns, and assembly) were repeated three additional times for a total of four blocks. We then counted 1) the number of pegs placed in the left column, 2) the number of pegs placed in the right column, 3) the number of pairs of pegs placed in a row, and 4) the number of items (i.e., peg, screw, or washer) appropriately placed on the pegboard.

During the Sequence Learning Task, participants were presented with four squares on a computer screen that were shaded to black one at a time. After a brief practice phase at half-speed, participants performed a test phase consisting of 6 random blocks, 18 trials each, where shaded squares were presented in a random order and 9 sequence blocks, 36 trials each, where shaded squares were presented in a 6-element pattern (i.e., 1-3-2-3-4-2 with 1 representing the leftmost box). Paralleling Kwak et al. (2012), we used the following block order: random-sequence-sequence-sequence-random, which was repeated 3 times. Given that the task was intended to be explicit, the nature of each upcoming block was disclosed to participants, who were also informed that their goal was to learn a sequence. Participants were instructed to respond to each black-shaded square with the corresponding key on the computer keyboard, using their left middle finger for the leftmost box and their left index, right index, and right middle fingers for the remaining boxes from left to right. Shaded squares were presented on the screen for 200 ms each, and participants were asked to respond as quickly and accurately as possible within 800 ms. One participant was excluded from calculations for this task as 98% of sequence trials were missed, suggesting that they did not adequately perform the task as instructed.

Z-scores for the Postural Sway Task were transformed (signs flipped) as greater sway area represents poorer balance. One participant was excluded from calculations for the Postural Sway Task due to possible technical errors, and one participant was excluded from the Sequence Learning Task for missing more than 90% of trials. As such, the motor composite for these individuals was calculated by averaging z-scores from only 2/3 motor tasks.

*A5. Sleep Assessments*

With the overnight actigraphy watches, sleep data was collected in 60-second epochs for 24 hours each day. To detect sleep periods, we used the default ActiGraph algorithm provided with the ActiLife software. The GT3X watches are equipped with a 3-axis accelerometer and wear time sensors, which collected data used to complete automatic wear time validation via the Troiano algorithm. Any non-wear periods were subsequently excluded from analysis. To score sleep periods, we used the Cole-Kripke algorithm, which is specifically tailored to adult populations (Cole et al., 1992).

Due to technical watch malfunctions, sleep data was not collected for 3 subjects. Further, due to battery issues, a full 10 days’ worth of data was not achieved for 8 subjects. Of these subjects, 6 had 9 days of data and 2 only had 4 days of data. However, sleep scoring was still carried out for these subjects by averaging across available days.

*A6. Neuroimaging*

For structural MRI, we used a high-resolution T1-weighted 3D magnetization prepared rapid gradient multi-echo (MPRAGE) scan (repetition time (TR) = 2400 ms; acquisition time = 7 minutes; voxel size = 0.8 mm^3^) and a high-resolution T2-weighted scan (TR = 3200 ms; acquisition time = 5.5 minutes; voxel size = 0.8 mm^3^), each with a multiband acceleration factor of 2.

For resting-state imaging, we administered four blood-oxygen level dependent (BOLD) functional connectivity (fcMRI) scans with the following parameters: multiband factor of 8, 488 volumes, TR of 720 ms, and 2.5 mm^3^ voxels. Each fcMRI scan was set to 6 minutes for a total of 24 minutes of resting-state imaging, and scans were acquired with alternating phase encoding directions (i.e., two anterior to posterior scans and two posterior to anterior scans). During fcMRI scans, participants were asked to lie still with their eyes open while fixating on a central cross.

Images were converted from DICOM to NIFTI and organized into a Brain Imaging Data Structure (BIDS, version 1.6.0) format via the latest docker container version of bidskit (version 2021.6.14, <https://github.com/jmtyszka/bidskit>). Using the split tool distributed with the FMRIB Software Library (FSL) package (Jenkinson et al., 2012), a single volume was extracted from two oppositely encoded BOLD images to estimate B_0_ field maps.

For the cortical ROIs, 3.5 mm spherical seeds were creating in FSL; the DLPFC seed was centered at (MNI: -36, 48, 10) and the M1 seed was centered at (MNI: -46, 1, 21) using the Johns Hopkins University (JHU) atlas (Lim et al., 2013). Both cortical seeds were placed in the left hemisphere.

Table A2. *Cortical Seed Origins.*

| *Region* | *Cluster Size* | *MNI Coordinates* | | | *P_(FDR)_* |
| --- | --- | --- | --- | --- | --- |
|  |  | *X* | *Y* | *Z* |  |
| *Right Crus I* | | | | | |
| Left Dorsolateral Prefrontal Cortex | 52 | -38 | 44 | 38 | 0.023 |
| *Right Lobule V* | | | | | |
| Left Primary Motor Cortex | 64 | -64 | -8 | 36 | 0.014 |

Seed-to-voxel CBLM whole-brain results from the prior work (Ballard et al., 2022) used to identify cortical seeds for the current work.

*A7. Exploratory Results*

Post-hoc t-tests between different pairs of reproductive stages for each hormone type signify significant differences between reproductive and early postmenopausal females for 17$\beta$-estradiol (*t*(28.99) = 2.87, *p* = 0.008) and progesterone (*t*(13.59) = 2.48, *p* = 0.027) as well as differences between reproductive and late postmenopausal females for both 17$\beta$-estradiol (*t*(25.89) = 3.84, *p* < 0.001) and progesterone (*t*(12.53) = 2.82, *p* = 0.015). No significant differences were found between any reproductive stage pairs for testosterone (all *ps* > 0.617). In addition, significant differences between early and late postmenopausal females were not observed across hormone types (all *ps* > 0.335). The resulting range for each sex hormone within our sample of healthy adult females is consistent with other work that incorporates salivary measures in similar groups (Felmingham et al., 2021; Gandara et al., 2007; Gavrilova and Lindau, 2009).

**References**

Cole, R.J., Kripke, D.F., Gruen, W., Mullaney, D.J., Gillin, J.C., 1992. Automatic sleep/wake identification from wrist activity. Sleep 15, 461–469. doi:10.1093/sleep/15.5.461

Felmingham, K.L., Caruana, J.M., Miller, L.N., Ney, L.J., Zuj, D.V., Hsu, C.M.K., Nicholson, E., To, A., Bryant, R.A., 2021. Lower estradiol predicts increased reinstatement of fear in women. Behav. Res. Ther. 142, 103875. doi:10.1016/j.brat.2021.103875

Gandara, B.K., Leresche, L., Mancl, L., 2007. Patterns of salivary estradiol and progesterone across the menstrual cycle. Ann. N. Y. Acad. Sci. 1098, 446–450. doi:10.1196/annals.1384.022

Gavrilova, N., Lindau, S.T., 2009. Salivary sex hormone measurement in a national, population-based study of older adults. J. Gerontol. B, Psychol. Sci. Soc. Sci. 64 Suppl 1, i94-105. doi:10.1093/geronb/gbn028

Jenkinson, M., Beckmann, C.F., Behrens, T.E., Woolrich, M.W., Smith, S.M., 2012. FSL. Neuroimage 62, 782–790. doi:10.1016/j.neuroimage.2011.09.015

Kwak, Y., Müller, M.L.T.M., Bohnen, N.I., Dayalu, P., Seidler, R.D., 2012. l-DOPA changes ventral striatum recruitment during motor sequence learning in Parkinson’s disease. Behav. Brain Res. 230, 116–124. doi:10.1016/j.bbr.2012.02.006

Lim, I.A.L., Faria, A.V., Li, X., Hsu, J.T.C., Airan, R.D., Mori, S., van Zijl, P.C.M., 2013. Human brain atlas for automated region of interest selection in quantitative susceptibility mapping: application to determine iron content in deep gray matter structures. Neuroimage 82, 449–469. doi:10.1016/j.neuroimage.2013.05.127
