## Appendix B for "Hormone-sleep interactions predict cerebellar connectivity and behavior in aging females"

Table B1. *Results from the Best Fit Model for Crus I-to-DLPFC Connectivity.*

| *Crus I-to-DLPFC Connectivity* | | | | |
| --- | --- | --- | --- | --- |
| *Predictor* | *Beta* | *SE* | *T* | *P* |
| *Fitted Model* |  |  |  |  |
| Intercept | 0.972 | 0.942 | 1.032 | 0.309 |
| Late Postmenopause | 0.162 | 0.035 | 4.645 | < 0.001*** |
| Reproductive | 0.056 | 0.043 | 1.297 | 0.203 |
| 17$\beta$-Estradiol | 1.213 | 0.799 | 1.518 | 0.138 |
| Progesterone | -0.011 | 0.006 | -1.731 | 0.092 |
| Sleep Quantity | -0.022 | 0.036 | -0.601 | 0.552 |
| Sleep Quality | -0.009 | 0.012 | -0.732 | 0.469 |
| 17$\beta$-Estradiol*Sleep Quantity | 0.110 | 0.042 | 2.623 | 0.013* |
| 17$\beta$-Estradiol*Sleep Quality | -0.023 | 0.010 | -2.253 | 0.030* |
| Progesterone*Sleep Quantity | -0.001 | 0.000 | -2.383 | 0.022* |
| Progesterone*Sleep Quality | 0.000 | 0.000 | 2.205 | 0.034* |
| *F*(10, 37) = 5.484*, R*^2^_adj_ = 0.488*, p* = 0.00006, *1-*$\beta$ = 0.99 | | | | |

Table B2. *Results from the Original Model for Crus I-to-DLPFC Connectivity.*

| *Crus I-to-DLPFC Connectivity* | | | | |
| --- | --- | --- | --- | --- |
| *Predictor* | *Beta* | *SE* | *T* | *P* |
| *Original Model* |  |  |  |  |
| Intercept | -0.182 | 3.656 | -0.050 | 0.960 |
| Late Postmenopause | 0.159 | 0.036 | 4.386 | < 0.001*** |
| Reproductive | 0.048 | 0.047 | 1.024 | 0.313 |
| 17$\beta$-Estradiol | 1.248 | 0.828 | 1.508 | 0.141 |
| Progesterone | -0.011 | 0.006 | -1.746 | 0.090 |
| Testosterone | 0.000 | 0.001 | -0.563 | 0.577 |
| Sleep Quantity | 0.161 | 0.508 | 0.316 | 0.754 |
| Sleep Quality | 0.005 | 0.042 | 0.113 | 0.911 |
| 17$\beta$-Estradiol*Sleep Quantity | 0.111 | 0.044 | 2.536 | 0.016* |
| 17$\beta$-Estradiol*Sleep Quality | -0.023 | 0.011 | -2.197 | 0.035* |
| Progesterone*Sleep Quantity | -0.001 | 0.000 | -2.391 | 0.022* |
| Progesterone*Sleep Quality | 0.000 | 0.000 | 2.219 | 0.033* |
| Sleep Quantity*Sleep Quality | -0.002 | 0.006 | -0.359 | 0.722 |
| *F*(12, 35) = 4.398*, R*^2^_adj_ = 0.465*, p* = 0.0003 | | | | |

Table B3. *Results from the Best Fit Model for Lobule V-to-M1 Connectivity.*

| *Lobule V-to-M1 Connectivity* | | | | |
| --- | --- | --- | --- | --- |
| *Predictor* | *Beta* | *SE* | *T* | *P* |
| *Fitted Model* |  |  |  |  |
| Intercept | 4.483 | 2.626 | 1.707 | 0.096 |
| Late Postmenopause | 0.039 | 0.028 | 1.401 | 0.169 |
| Reproductive | -0.025 | 0.036 | -0.686 | 0.497 |
| 17$\beta$-Estradiol | 0.399 | 0.160 | 2.489 | 0.017* |
| Progesterone | -0.007 | 0.004 | -1.736 | 0.091 |
| Sleep Quantity | -0.620 | 0.381 | -1.629 | 0.112 |
| Sleep Quality | -0.538 | 0.030 | -1.807 | 0.079 |
| 17$\beta$-Estradiol*Sleep Quantity | -0.062 | 0.025 | -2.483 | 0.018* |
| Progesterone*Sleep Quality | 0.000 | 0.000 | 1.778 | 0.084 |
| Sleep Quantity*Sleep Quality | 0.007 | 0.004 | 1.740 | 0.090 |
| *F*(9, 38) =1.843 *, R*^2^_adj_ = 0.139*, p* = 0.092, *1-*$\beta$ = 0.53 | | | | |

Table B4. *Results from the Original Model for Lobule V-to-M1 Connectivity.*

| *Lobule V-to-M1 Connectivity* | | | | |
| --- | --- | --- | --- | --- |
| *Predictor* | *Beta* | *SE* | *T* | *P* |
| *Original Model* |  |  |  |  |
| Intercept | 2.881 | 2.940 | 0.980 | 0.334 |
| Late Postmenopause | 0.035 | 0.029 | 1.185 | 0.244 |
| Reproductive | -0.035 | 0.038 | -0.929 | 0.359 |
| 17$\beta$-Estradiol | 1.067 | 0.666 | 1.603 | 0.118 |
| Progesterone | -0.011 | 0.005 | -2.052 | 0.048* |
| Testosterone | 0.000 | 0.000 | -0.818 | 0.419 |
| Sleep Quantity | -0.462 | 0.409 | -1.130 | 0.266 |
| Sleep Quality | -0.035 | 0.034 | -1.027 | 0.311 |
| 17$\beta$-Estradiol*Sleep Quantity | -0.044 | 0.035 | -1.240 | 0.223 |
| 17$\beta$-Estradiol*Sleep Quality | -0.009 | 0.009 | -1.032 | 0.309 |
| Progesterone*Sleep Quantity | 0.000 | 0.000 | -0.431 | 0.669 |
| Progesterone*Sleep Quality | 0.000 | 0.000 | 1.917 | 0.063 |
| Sleep Quantity*Sleep Quality | 0.006 | 0.005 | 1.194 | 0.241 |
| *F*(12, 35) = 1.482*, R*^2^_adj_ = 0.110*, p* = 0.178 | | | | |

Table B5. *Results from the Best Fit Model for Cognitive Performance.*

| *Cognitive Performance* | | | | |
| --- | --- | --- | --- | --- |
| *Predictor* | *Beta* | *SE* | *T* | *P* |
| *Fitted Model* |  |  |  |  |
| Intercept | -30.050 | 19.370 | -1.551 | 0.129 |
| 17$\beta$-Estradiol | -0.639 | 5.217 | -0.122 | 0.903 |
| Progesterone | -0.055 | 0.048 | -1.159 | 0.253 |
| Testosterone | -0.010 | 0.003 | -3.010 | 0.005** |
| Sleep Quantity | 5.325 | 2.767 | 1.924 | 0.062 |
| Sleep Quality | 0.367 | 0.220 | 1.673 | 0.102 |
| Crus I-to-DLPFC Connectivity | 0.741 | 7.257 | 0.102 | 0.919 |
| Lobule V-to-M1 Connectivity | 48.870 | 26.750 | 1.827 | 0.075 |
| 17$\beta$-Estradiol*Sleep Quantity | 0.518 | 0.236 | 2.199 | 0.034* |
| 17$\beta$-Estradiol*Sleep Quality | -0.029 | 0.067 | -0.431 | 0.669 |
| Progesterone*Sleep Quantity | -0.004 | 0.002 | -2.186 | 0.035* |
| Progesterone*Sleep Quality | 0.001 | 0.001 | 1.586 | 0.121 |
| Sleep Quantity*Sleep Quality | -0.063 | 0.031 | -2.050 | 0.047* |
| 17$\beta$-Estradiol*Crus I-to-DLPFC | 1.691 | 1.216 | 1.391 | 0.172 |
| 17$\beta$-Estradiol*Lobule V-to-M1 | -2.472 | 1.775 | -1.393 | 0.172 |
| Progesterone*Crus I-to-DLPFC | 0.004 | 0.012 | 0.361 | 0.720 |
| Sleep Quantity*Crus I-to-DLPFC | -0.768 | 1.037 | -0.740 | 0.463 |
| Sleep Quantity*Lobule V-to-M1 | -1.834 | 1.179 | -1.555 | 0.128 |
| Sleep Quality*Lobule V-to-M1 | -0.380 | 0.313 | -1.215 | 0.232 |
| *F*(18, 39) = 3.895*, R*^2^_adj_ = 0.478*, p* = 0.0002, *1-*$\beta$ = 0.99 | | | | |

Table B6. *Results from the Original Model for Cognitive Performance.*

| *Cognitive Performance* | | | | | | | |
| --- | --- | --- | --- | --- | --- | --- | --- |
| *Predictor* | *Beta* | | *SE* | *T* | | | *P* |
| *Original Model* |  | |  |  | | |  |
| Intercept | -33.000 | | 25.050 | -1.317 | | | 0.200 |
| Late Postmenopause | -0.045 | | 0.310 | -0.145 | | | 0.886 |
| Reproductive | -0.224 | | 0.322 | -0.694 | | | 0.494 |
| 17$\beta$-Estradiol | -4.218 | | 7.118 | -0.593 | | | 0.559 |
| Progesterone | -0.073 | | 0.073 | -0.999 | | | 0.327 |
| Testosterone | -0.010 | | 0.004 | -2.428 | | | 0.023* |
| Sleep Quantity | 7.312 | | 3.494 | 2.093 | | | 0.047* |
| Sleep Quality | 0.371 | | 0.288 | 1.288 | | | 0.210 |
| Crus I-to-DLPFC Connectivity | -14.740 | 25.450 | | | -0.579 | 0.568 | |
| Lobule V-to-M1 Connectivity | 62.970 | 34.900 | | | 1.804 | 0.083 | |
| 17$\beta$-Estradiol*Sleep Quantity | 0.256 | 0.376 | | | 0.683 | 0.501 | |
| 17$\beta$-Estradiol*Sleep Quality | 0.032 | 0.095 | | | 0.341 | 0.736 | |
| Progesterone*Sleep Quantity | -0.005 | 0.003 | | | -1.747 | 0.093 | |
| Progesterone*Sleep Quality | 0.001 | 0.001 | | | 1.296 | 0.207 | |
| Sleep Quantity*Sleep Quality | -0.081 | 0.039 | | | -2.092 | 0.047* | |
| 17$\beta$-Estradiol*Crus I-to-DLPFC | 2.143 | 1.586 | | | 1.351 | 0.189 | |
| 17$\beta$-Estradiol*Lobule V-to-M1 | -4.644 | 3.621 | | | -1.283 | 0.211 | |
| Progesterone*Crus I-to-DLPFC | 0.004 | 0.017 | | | 0.239 | 0.813 | |
| Progesterone*Lobule V-to-M1 | -0.001 | 0.056 | | | -0.015 | 0.988 | |
| Sleep Quantity*Crus I-to-DLPFC | -2.211 | 1.491 | | | -1.483 | 0.151 | |
| Sleep Quality*Crus I-to-DLPFC | 0.281 | 0.307 | | | 0.913 | 0.370 | |
| Sleep Quantity*Lobule V-to-M1 | -1.086 | 1.594 | | | -0.681 | 0.502 | |
| Sleep Quality*Lobule V-to-M1 | -0.572 | 0.414 | | | -1.381 | 0.180 | |
| *F*(22, 25) = 2.668*, R*^2^_adj_ = 0.439*, p* = 0.0097 | | | | | | | |

Table B7. *Results from the Best Fit Model for Motor Performance.*

| *Motor Performance* | | | | |
| --- | --- | --- | --- | --- |
| *Predictor* | *Beta* | *SE* | *T* | *P* |
| *Fitted Model* |  |  |  |  |
| Intercept | -43.348 | 15.290 | -2.835 | 0.006** |
| 17$\beta$-Estradiol | 8.013 | 3.398 | 2.358 | 0.021* |
| Testosterone | -0.008 | 0.002 | -3.274 | 0.002** |
| Sleep Quantity | 4.660 | 2.182 | 2.135 | 0.037* |
| Sleep Quality | 0.510 | 0.175 | 2.908 | 0.005** |
| 17$\beta$-Estradiol*Sleep Quantity | 0.304 | 0.134 | 2.272 | 0.026* |
| 17$\beta$-Estradiol*Sleep Quality | -0.107 | 0.043 | -2.503 | 0.015* |
| Sleep Quantity*Sleep Quality | -0.056 | 0.025 | -2.260 | 0.027* |
| *F*(7, 65) = 4.159*, R*^2^_adj_ = 0.235*, p* = 0.0008, *1-*$\beta$ = 0.94 | | | | |

Table B8. *Results from the Original Model for Motor Performance.*

| *Motor Performance* | | | | |
| --- | --- | --- | --- | --- |
| *Predictor* | *Beta* | *SE* | *T* | *P* |
| *Original Model* |  |  |  |  |
| Intercept | -30.100 | 20.210 | -1.489 | 0.149 |
| Late Postmenopause | -0.184 | 0.251 | -0.733 | 0.471 |
| Reproductive | 0.203 | 0.260 | 0.779 | 0.443 |
| 17$\beta$-Estradiol | 8.388 | 5.743 | 1.461 | 0.157 |
| Progesterone | -0.023 | 0.059 | -0.381 | 0.706 |
| Testosterone | -0.006 | 0.003 | -1.742 | 0.094 |
| Sleep Quantity | 2.994 | 2.819 | 1.062 | 0.298 |
| Sleep Quality | 0.351 | 0.232 | 1.510 | 0.144 |
| Crus I-to-DLPFC Connectivity | -12.640 | 20.530 | -0.615 | 0.544 |
| Lobule V-to-M1 Connectivity | 0.712 | 28.160 | 0.025 | 0.980 |
| 17$\beta$-Estradiol*Sleep Quantity | 0.355 | 0.303 | 1.171 | 0.253 |
| 17$\beta$-Estradiol*Sleep Quality | -0.117 | 0.077 | -1.520 | 0.141 |
| Progesterone*Sleep Quantity | 0.000 | 0.002 | -0.170 | 0.866 |
| Progesterone*Sleep Quality | 0.000 | 0.001 | 0.357 | 0.724 |
| Sleep Quantity*Sleep Quality | -0.035 | 0.031 | -1.120 | 0.274 |
| 17$\beta$-Estradiol*Crus I-to-DLPFC | -1.703 | 1.280 | -1.331 | 0.195 |
| 17$\beta$-Estradiol*Lobule V-to-M1 | 4.474 | 2.922 | 1.531 | 0.138 |
| Progesterone*Crus I-to-DLPFC | 0.017 | 0.014 | 1.213 | 0.236 |
| Progesterone*Lobule V-to-M1 | 0.024 | 0.045 | 0.537 | 0.596 |
| Sleep Quantity*Crus I-to-DLPFC | -0.881 | 1.203 | -0.732 | 0.471 |
| Sleep Quality*Crus I-to-DLPFC | 0.211 | 0.248 | 0.851 | 0.403 |
| Sleep Quantity*Lobule V-to-M1 | 0.340 | 1.286 | 0.264 | 0.794 |
| Sleep Quality*Lobule V-to-M1 | -0.111 | 0.334 | -0.333 | 0.742 |
| *F*(22, 25) = 2.538*, R*^2^_adj_ = 0.419*, p* = 0.0132 | | | | |

**References**

Ballard, H.K., Jackson, T.B., Hicks, T.H., Bernard, J.A., 2022. The association of reproductive stage with lobular cerebellar network connectivity across female adulthood. Neurobiol. Aging 117, 139–150. doi:10.1016/j.neurobiolaging.2022.05.014
